# Identification and determination of the urinary metabolite of iodotyrosine *in vivo*

**DOI:** 10.1101/2024.09.30.615905

**Authors:** Cindy Xinyu Ji, Danae Zonias, Nafisa Djumaeva, Ranchu Cheng, Kawther Salim, Pouya Alikhani, Tope Oyelade, Kevin P. Moore, José C. Moreno, Ali R. Mani

## Abstract

**Background:** Congenital hypothyroidism screening traditionally relies on detecting elevated thyroid-stimulating hormone levels, yet this approach may not detect a specific type of congenital hypothyroidism caused by iodotyrosine dehalogenase-1 (*Dehal1*) deficiency. The deficiency of this enzyme prevents the deiodination of mono-iodotyrosine (MIT) and di-iodotyrosine (DIT) in the process of iodine recycling. This underscores the potential use of iodotyrosine or its metabolites as non-invasive urinary biomarkers for early diagnosis of congenital hypothyroidism. However, the urinary metabolites of MIT/DIT have not yet been discovered. Thus, this study aimed to identify the urinary metabolites of iodotyrosine in experimental models.

**Method:** Gas chromatography mass spectrometry was used to identify the urinary metabolites of iodotyrosine following intraperitoneal injection of MIT in rats. An isotope dilution mass spectrometric assay was developed for assessment of identified metabolites. Urine samples from *Dehal1* knockout mice were used to confirm the results.

**Results:** We identified novel iodotyrosine metabolites, 3-iodo-4-hydroxyphenylacetic acid (IHPA), and 3,5-diiodo-4-hydroxyphenylacetic acid (Di-IHPA) as the primary urinary metabolites of MIT and DIT respectively. The concentrations of urinary IHPA and Di-IHPA were significantly higher in *Dehal1* knockout mice.

**Conclusion:** Our findings suggest that IHPA is detected in larger quantities and may hold more clinical significance than previously identified biomarkers like MIT and DIT, making it a promising candidate for diagnosing congenital hypothyroidism or other conditions associated with iodine recycling inhibition.

## Introduction

Iodine is an essential element actively taken up by the thyroid gland for the synthesis of thyroid hormones. Free mono-iodotyrosines (MIT) and di-iodotyrosine (DIT) are important by-products in the process of thyroid hormone synthesis. In healthy individuals, to prevent the loss of iodine in the urine as iodotyrosines, iodine in MIT and DIT is recycled in the thyroid gland. This recycling process is catalysed by the enzyme iodotyrosine deiodinase, also known as iodotyrosine dehalogenase-1 (DEHAL1) [1]. Notably, this enzyme is not only expressed in the thyroid but is also present in the kidney, liver, and choroid plexus, underscoring the importance of iodine homeostasis in different organ systems [2].

One significant consequence of the intrathyroidal recycling of iodide derived from the dehalogenation of MIT/DIT is the preservation of iodine that has not been incorporated into the thyroid hormone structure. Indeed, patients with congenital dehalogenase-1 deficiency may develop hypothyroidism, underscoring the importance of this recycling mechanism in health and disease [3-5]. The significance of this intricate interplay was further emphasized by the implication of its mutations in various cases of congenital hypothyroidism in 2008 [3,6]. More recently, the observation of preclinical hypothyroidism in a knockout mice model of dehalogenase-1 deficiency has strengthened the evidence for the role of this enzyme in normal thyroid physiology [2]. The primary anomaly observed in such cases is a disruption in both intrathyroidal and peripheral deiodination of iodotyrosines, leading to the loss of iodinated organic compounds in the urine [2,7].

Despite the crucial role of dehalogenase in iodine recycling, current tools for identifying congenital hypothyroidism are unable to detect deficiency of this enzyme [8]. The current screening methods for congenital hypothyroidism primarily detect elevated thyroid-stimulating hormone (TSH) [9]. However, this method cannot easily detect congenital hypothyroidism due to dehalogenase-1 deficiency. Therefore, given the importance of iodine in neural development [10], it is crucial to find a more effective biomarker to detect the condition before any potential neural developmental complications occur.

Since iodinated tyrosine can be excreted in the urine, a recent study has proposed an alternative method by focusing on the detection of MIT and DIT in urine. However, the presented concentrations are on a scale of 10^-2^ ng per g of urine, and DIT is nearly undetectable in most of the samples [11]. Furthermore, recent studies report that excreted MIT/DIT can only account for a small fraction of the total amount of iodinated compounds lost in the urine [2]. These findings suggest the possibility of another pathway through which iodine loss occurs, likely attributed to the metabolites of iodotyrosine.

Tyrosine undergoes metabolism in the liver and other peripheral tissues, typically involving deamination and decarboxylation to form 4-hydroxyphenyl acetate (4-HPA). Previous reports from our group have indicated that halogenated derivatives of tyrosine, such as 3-chlorotyrosine and 3-bromotyrosine, are excreted in the urine as 3-chloro-4-hydroxyphenyl acetate (CHPA) and 3-bromo-4-hydroxyphenyl acetate (BHPA), with urinary concentrations significantly higher than their unmetabolized forms [12-14]. The metabolic pathway for MIT and DIT is not well understood. In addition to dehalogenation, these compounds can undergo further metabolism in the liver. Therefore, it is crucial to identify the main urinary metabolite of iodotyrosines *in vivo*. This identification could be instrumental in developing novel methods for the early detection of dehalogenase-1 deficiency. Building on this groundwork, the current study aims to identify iodotyrosine metabolites in urine.

## Method

### 1. Ethics statement

Animal studies followed the guidelines from the European Animal Research Ethical Committee and Home Office UK and were approved by the Animal Research Ethical Committee of the Autonomous University of Madrid, Spain (Proex-13/17 and Proex-141.1/23).

### 2. Identification of the urinary metabolites of MIT in rats

To identify the urinary metabolite of MIT *in vivo*, we injected 30 µg of MIT solution in isotonic phosphate buffer saline (PBS) or the vehicle alone (PBS) intraperitoneally to 3 healthy Sprague-Dawley rats. The rats had access to food and water *ad libitum* and were kept in standard light/dark cycle. 24 hours urine samples were collected using metabolic cages. All experiments were carried out under the Home Office (UK) licence. Collected urines were stored at -80°C until analysis.

### 3. Synthesis 3-Iodo-4-HydroxyPhenylacetic Acid (IHPA)

Iodo-hydroxyphenylacetic acid was synthesised *in vitro* by a redox reaction that oxidises iodide in the presence of 4-hydroxyphneylacetic acid (4-HPA). In brief, 0.2 ml of sodium iodide (30 mg/ml), 0.2 ml of potassium iodate (20 mg/ml) and 0.2 ml of HPA (30 mg/ml) were mixed in a plastic tube and were then mixed with 0.2 ml of HCl (1 M). The mixture was centrifuged three times (once at 1,000 x g, for 2 minutes then twice at 10,000 x g, for 5 minutes each) followed by washing twice with 1 ml of distilled water. The resulting cloudy white part of the solution was collected and dried down under stream of nitrogen.

### 4. HPLC for separation of iodinated derivatives of 4-hydroxyphneylacetic acid

Liquid chromatography was conducted using a C18 column on a Jasco HPLC machine connected to a photodiode array UV detector (Jasco Co., Tokyo, Japan). A gradient of two mobile phase solutions (A and B) were used to separate molecules by their difference in polarity and size. Solution A was made of 10% acetonitrile in ultrapure water with 0.08% of trifluoroacetic acid. Solution B was made of 70% acetonitrile in ultrapure water with 0.08% of trifluoroacetic acid. The flow rate of the mobile phase was set to 0.4 ml/min with injection volume of 10 μl of sample in each run. The process initiated with 100% of solution A and 0% of solution B for one minute. Then the proportion of solution B gradually increased to 80% in 25 minutes. Solution B stayed at 80% for further 5 min before returning to 0% in 5 min. Considering the phenolic structure of 4-HPA and its derivatives, the chromatogram was selected for viewing at a wavelength of 220 nm.

### 5. Synthesis of ^13^C-labelled 3-Iodo-4-hydroxyphenylacetic Acid (IHPA)

^13^C-labelled IHPA was synthesised by iodination of ^13^C-labelled 4-HPA as described above. ^13^C-4-HPA is not commercially available, thus we synthesized it via deamination and decarboxylation of ^13^C_9_-tyrosine using a snake venom enzyme procedure that has been described before [15]. It was purified using a 0.22 µm pore filter and 1.5 ml of ethyl acetate to extract the compound. The products were further purified using C18 solid phase extraction. It was then followed by HPLC, collecting the liquid at the retention time of 17.6 min as described above and drying under nitrogen.

### 6. Confirmation and detection by GC/MS using full scan mass spectrometry.

#### 6.1. Synthetic IHPA

IHPA was derivatised to its pentafluorobenzyl ester using 40 μl of 10% pentafluorobenzyl bromide (PFBBr) in acetonitrile and 20 μl of 10% diisopropylethylamine (DIPEA) in acetonitrile as a catalyst. After 30 minutes incubation at room temperature, the solution was dried under nitrogen, resuspended in 50 μl hexane and then injected into the GC/MS [12,13,15].

Samples were analysed in negative-ion chemical ionization mode with methane as the reagent gas and helium gas as the carrier gas. The carrier gas flow was maintained at 1 ml/min with 1 µl of each sample injected for analysis. The initial column temperature was maintained at 150 °C for 1 minute and then increased to 300 °C at 20 °C/min. The ion source and interface temperatures were set at 200 and 320 °C respectively, and a full scan was performed (100-1000 m/z). An Agilent instrument (GCMS 5977 Turbo System, Agilent Technologies, Palo Alto, CA, USA) was used for gas chromatography mass spectrometry.

#### 6.2. Identification of MIT urinary metabolites in rat urine

20 µl of urine sample from control or MIT-treated rat urine was used. For buffering, 100 µl of citrate buffer (0.1 M, pH=5.0) was added to the sample, followed by 1 ml of ethyl acetate for organic solvent extraction. After vortex mixing, 500 µl of supernatant (ethyl acetate phase) was dried down using nitrogen. The dried precipitates were derivatized using 20 µl of 10% DIPEA and 40 µl of 10% PFBBr for 60 min as described. This was then dried down under nitrogen and was redissolved in 50 µl of hexane before injection into GC/MS for full scan (100-1000 m/z).

### 7. Detection of IHPA and Di-IHPA in mice urine using single ion monitoring (SIM) with ^13^C-labelled internal standard.

Urine samples from *Dehal1* knockout mice were used in this section [2]. 20 µl urine was mixed with 2.5 ng of ^13^C-labeled IHPA and 100 µl of citrate buffer (0.1 M, pH=5.5). IHPA was extracted using organic solvent extraction by adding 1 ml ethyl acetate and drying the supernatant under nitrogen. The extracted samples were derivatised to a pentafluorobenzyl derivative by adding 20 µl of 10% DIPEA and 40 µl of 10% PFBBr for 1 h. This was then dried down under nitrogen and was redissolved in 50 µl of hexane before injection into GC/MS to perform a single ion monitoring mass spectrometry. Ions were monitored at m/z 456.9, and m/z 582.8 mass units for authentic IHPA and Di-IHPA respectively. We also monitored m/z 464.9 and m/z 590.8 for detection of ^13^C-labeled IHPA and Di-IHPA respectively.

Since the urinary concentration of metabolites can be affected by urine dilution, all results of the mice urine study were normalised for the urine specific gravity (SG) to eliminate the effect of hydration. To measure urinary specific gravity a digital urine refractometer (UG1, ATAGO co. Ltd, Tokyo, Japan) was used according to the manufacturer’s instructions. Data normalised using urine specific gravity by according to the formula by Harps et al., 2023 [16], where 1.025 is the standard urine specific gravity:

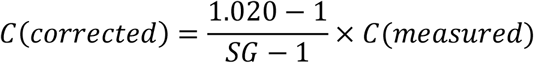

Mann-Whitney U test was used for statistical analysis in STATA (STATA/18.0, Stata Corp, TX, USA).

## Results

### 1. Full scan for identification and confirmation

#### 1.1 Structural confirmation of synthetic IHPA

Iodination of 4-HPA was associated with formation of two new peaks in the HPLC chromatogram with retention times of ∼17.6 and ∼23.4 min. The first peak had a UV absorption spectrum with maximum absorptions at 220 nm and 286 nm. The peak exhibited a UV absorption spectrum with maximum absorption at 220 nm, 240 and 286 nm. Following the derivatisation and GC/MS, two peaks were observed corresponding to two expected iodinated-HPA compounds, mono-iodo-HPA (IHPA) and di-iodo-HPA (Di-IHPA) (Figure 1). Analysis of the pentafluorobenzyl (PFB) ester of synthetic IHPA by full scan mode showed that IHPA has a bigger dominant ion at m/z 457 (retention time at 8.67min), which corresponds to the loss of a PFB group. Similarly, a fragment with m/z 582.8 (at retention time at 10.81 min) showed loss of PFB in Di-IHPA as shown in Figure 1. The full scan spectrum for ^13^C-labelled IHPA is shown in Appendix 1. The prominent ion was eight mass units heavier (i.e., m/z 465) than authentic IHPA (m/z 457) exhibiting the number of ^13^C atoms in the structure (^13^C_8_ IHPA). Based on these results, further work on urinary metabolites from *Dehal1*KO mice was focused on single ion monitoring at 457, 465 and 583 for IHPA, ^13^C_8_-IHPA and Di-IHPA respectively.

**Figure 1.**
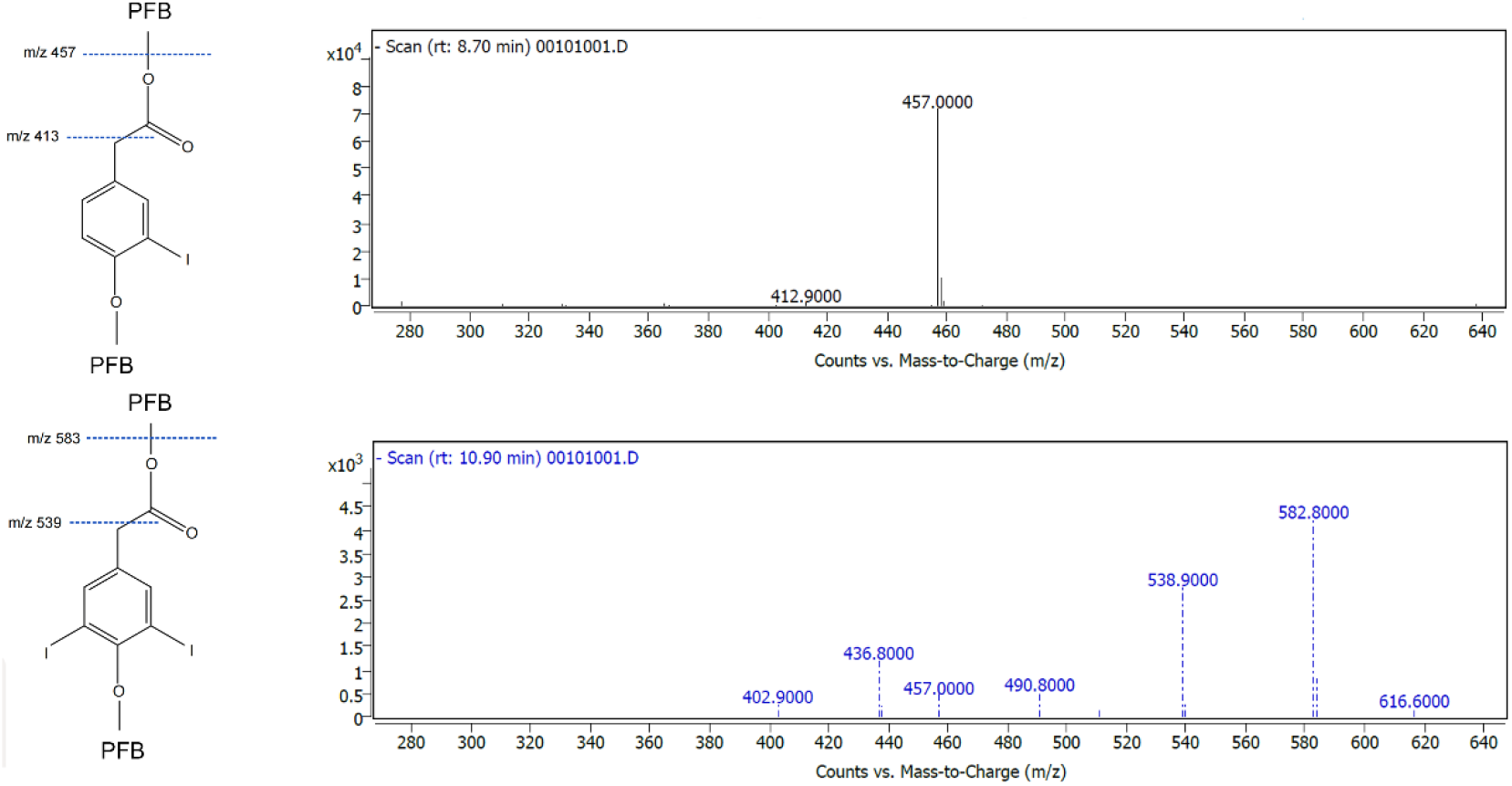
Structures and mass spectra from negative-ion chemical ionization scans of the pentafluorobenzyl derivative of authentic IHPA (upper panel) and Di-IHPA (lower panel).

#### 1.2 Metabolism of iodotyrosine by using rat urine 24 h after injecting MIT

As shown in Figure 2, analysis of urine extracts from rats that were administered MIT showed three main additional peaks on the gas chromatogram, one of which is at m/z 456.8 with a retention time of 8.68, which is identical to the synthetic peak of IHPA. In addition, two additional peaks of potential significance were observed at m/z 473, retention time of 9.2 and m/z 487 at a retention time of 9.93, respectively.

**Figure 2.**
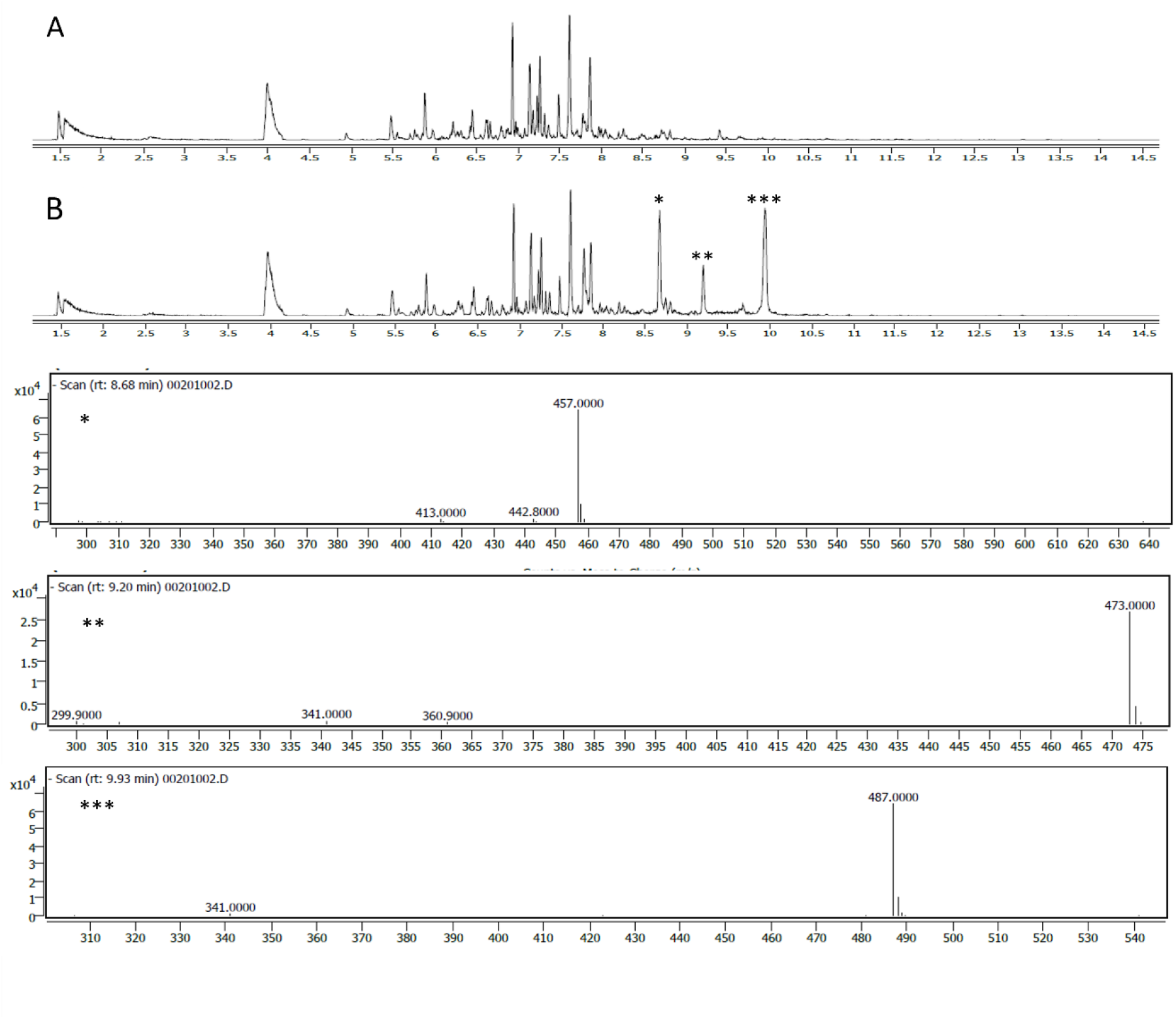
Gas chromatogram of urine samples analysed with full scan mass spectrometry after derivatization with pentafluorobenzyl bromide. The analysis of urine samples from rats given iodotyrosine (B) showed three consistent additional peaks on the chromatogram compared with urine samples of saline treated rats (A). The mass spectra of the first additional peak (*) is identical to the mass spectra of 3-iodo-4-hydroxyphenyl acetic acid (IHPA).

### 2 Detection and analysis of iodotyrosine metabolites in Dehal1KO mice urine using single ion monitoring (SIM).

^13^C-labelled IHPA was added to the urine samples collected from *Dehal1KO* mice. Following extraction and derivatisation, GC/MS with SIM was used with a focus on m/z at 457 and 583 for IHPA and Di-IHPA respectively. ^13^C-labelled mono-IHPA was monitored at m/z 465, which was 8 mass units heavier than the authentic IHPA (supplementary material 1). IHPA was detected in both wild-type and KO urine samples. However, Di-IHPA was detectable only in *Dehal1*KO samples (4 out of 5) and none of the wild-type samples. Box whisker plot for comparison of wild type and *Dehal1*KO are shown in Figure 3. Statistical analysis showed that the increase in IHPA and Di-IHPA were significant in *Dehal1*KO mice in comparison with control with p-values of 0.04 and 0.0073 respectively.

**Figure 3.**
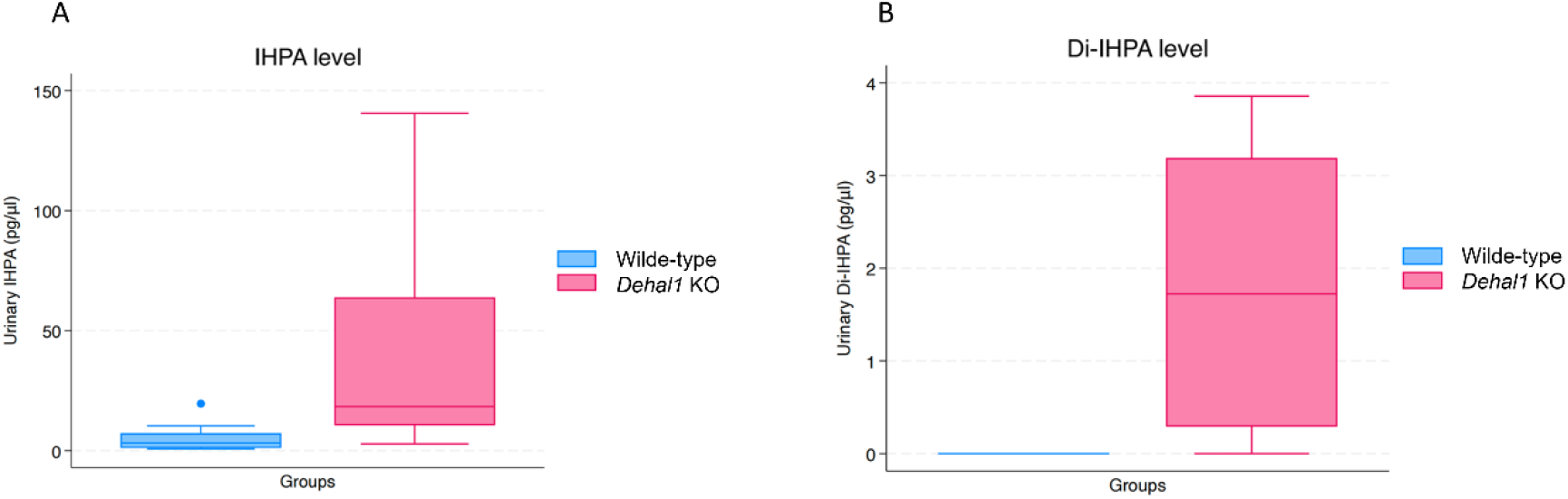
(A) Box whisker plot of IHPA data shows the level of IHPA is significantly increased in Dehal1KO mice compared with the control group. (B) Box whisker plot of Di-IHPA data shows Di-IHPA is exclusively detectable in the *Dehal1*KO group. However, the observed amount is notably smaller in scale when compared to IHPA.

## Discussion

In this study, our primary goal was to identify the urinary metabolites of MIT *in vivo*, aiming to discover a novel biomarker for the early diagnosis of dehalogenase-1 deficiency. Our results revealed the presence of at least one iodinated metabolite of MIT in experimental animals. To quantify this metabolite in mouse urine samples, we developed a sensitive tool using isotope dilution mass spectrometry.

Urinary analysis of rats administered MIT indicated the presence of IHPA as one of the metabolites. The metabolite profile of iodotyrosine in urine matched the chromatographic pattern on GC/MS with in vitro synthesized IHPA (m/z 457). As IHPA is not commercially available, we devised a simple protocol for its synthesis and purification, facilitating its use in future studies exploring its biochemical or physiological characteristics and its potential as a biomarker.

While our focus was on IHPA, the detection of two additional peaks suggests the potential existence of other metabolites with physiological significance, serving as potential biomarkers. Although we did not identify these metabolites in this study, the peak at m/z 473, 16 mass units heavier than iodo-HPA, raises the possibility of an additional oxygen atom, possibly indicating a compound like 3-iodo-4-hydroxymandelic acid. Further investigation is necessary to confirm the structure of these additional peaks.

In the *Dehal1*KO mice study, we utilized ^13^C-labelled IHPA as an internal standard at a known concentration, enhancing specificity and quantification of authentic metabolites in mouse urine. This methodology may be valuable in providing reference evidence for establishing diagnostic thresholds when using IHPA as a biomarker in the future. Considering the contribution of dehalogenase-1 in congenital dehalogenase deficiency and the observation that only a small proportion of iodinated tyrosine was lost in the form of intact MIT and DIT, on the scale of 10^-2^ ng per g [11], our study aimed to highlight additional urinary metabolites. Although Di-IHPA was undetectable in the wild type and became detectable in the KO mice, the difference in IHPA concentration was more pronounced, suggesting that IHPA might be a better potential biomarker due to its greater accessibility in the samples. Data were further analysed by converting the unit to ng per g of urine using the formula:

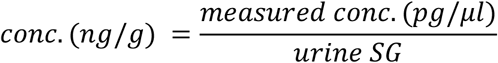

This was done to facilitate a direct comparison with previous findings regarding MIT and DIT. The results revealed a substantial presence of additional urinary metabolites, with a mean of 86.18 ng per g and 3.28 ng per g of IHPA and Di-IHPA, respectively. These quantities were found to be greater than those reported for MIT and DIT [2], emphasizing the significance of using IHPA and Di-IHPA as potential metabolite biomarkers in our study. The higher concentrations of urinary metabolites (IHPA and Di-IHPA), compared to urinary levels of MIT and DIT, may also explain why total iodine loss in the urine is significantly greater than the combined concentrations of unmetabolized MIT and DIT in dehalogenase-1 deficiency [2]. Future studies could focus on developing an assay that includes all urinary metabolites of iodotyrosines, enabling a more comprehensive assessment of patients’ iodine balance in health and disease.

The present study aligns with previous reports on the urinary metabolites of halogenated tyrosine. Deamination and decarboxylation of tyrosine are well-documented processes involving various carboxylases and monoamine oxidases. 3-nitrotyrosine, 3-chloro-tyrosine, and 3-bromotyrosine also undergo the same metabolic pathways [12-15,17]. There is potential to develop point-of-care devices for the measurement of urinary IHPA in the future. This aim is clinically significant because patients with congenital hypothyroidism due to dehalogenase-1 deficiency are typically not diagnosed early through routine screening tests (e.g., TSH), which increases their risk of neurodevelopmental deficits. Although we were able to detect IHPA in human urine (data not shown), we did not study the details of this metabolic pathway in humans in the present report. This limitation of the current study highlights the need for further research, and the results presented here may facilitate future investigations into the diagnostic and mechanistic potential of these iodinated biomolecules.

## Conclusion

In this study IHPA was identified as a urinary metabolite of MIT and a protocol was devised for the synthesis, purification and measurement of IHPA for future studies. Our findings indicate that IHPA is detected in larger quantities than previously identified biomarkers like MIT and DIT, making it a promising candidate for early diagnosing conditions associated with iodine recycling inhibition.

## Data Statement

The data that support the findings of this study are available on request from the corresponding author

## Authors contribution

The conception and design of the study (ARM, JCM, KPM), acquisition of data (CXJ, DZ, ND, RC, KS, PA, TO, ARM), analysis and interpretation of data (CXJ, DZ, ARM), drafted manuscript (CXJ), edited and revised manuscript (TO, PA, JCM, ARM), approved final version of manuscript (All authors).

---

**Supplementary material 1.**
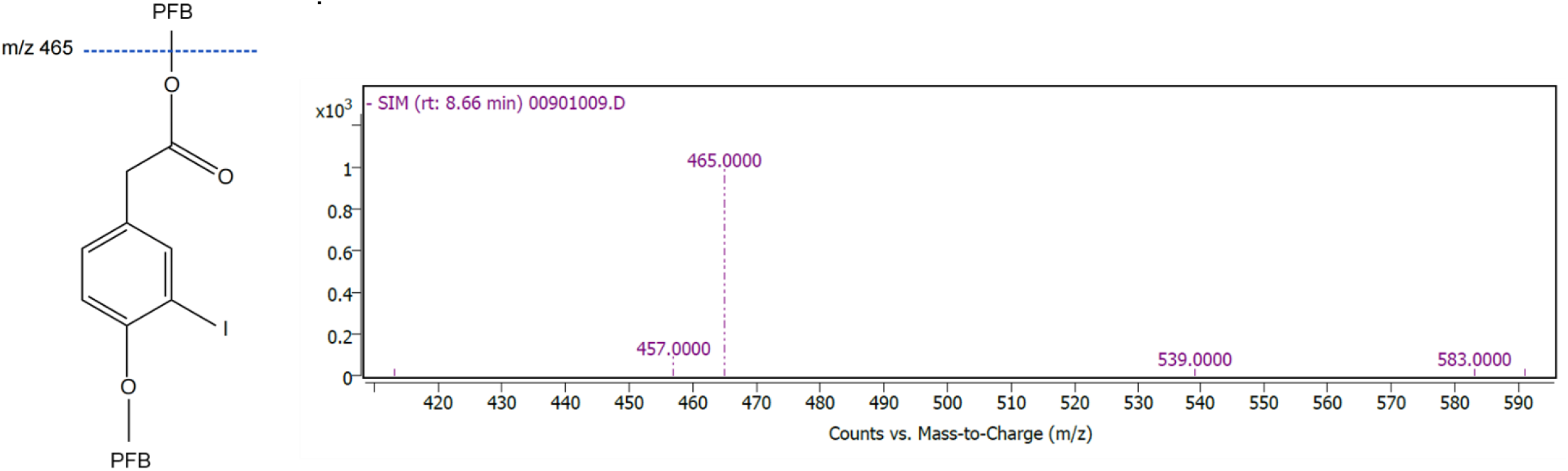
Structures and mass spectra from negative-ion chemical ionization scans of the pentafluorobenzyl derivative of ^13^C-lableled IHPA.

